# Dynamic network properties of the interictal brain determine whether seizures appear focal or generalised

**DOI:** 10.1101/576785

**Authors:** Wessel Woldman, Helmut Schmidt, Eugenio Abela, Fahmida A. Chowdhury, Adam D. Pawley, Sharon Jewell, Mark P. Richardson, John R. Terry

**Author notes:** **Corresponding author:** Wessel Woldman PhD, Living Systems Institute, Centre for Biomedical Modelling and Analysis, EPSRC Centre for Predictive Modelling in Healthcare, University of Exeter, UK. These authors made an equal contribution as last author (Mark P. Richardson, John R. Terry).

## Abstract

**Objective:** Current explanatory concepts suggest seizures emerge from ongoing dynamics of brain networks. It is unclear how brain network properties determine focal or generalised seizure onset, or how network properties can be described in a clinically-useful manner. Understanding network properties would cast light on seizure-generating mechanisms and allow to quantify in the clinic the extent to which a seizure is focal or generalised.

**Methods:** 68 people with epilepsy and 38 healthy controls underwent 19 channel scalp EEG recording. Functional brain networks were estimated in each subject using phase-locking between EEG channels in the 6-9Hz band from segments of 20s without interictal discharges. Simplified brain dynamics were simulated using a computer model. We introduce three concepts: Critical Coupling (C_c_), the ability of a network to generate seizures; Onset Index (OI), the tendency of a region to generate seizures; and Participation Index (PI), the tendency of a region to become involved in seizures.

**Results:** C_c_ was lower in both patient groups compared with controls. OI and PI were more variable in focal-onset than generalised-onset cases. No regions showed higher OI and PI in generalised-onset cases than in healthy controls; in focal cases, the regions with highest OI and PI corresponded to the side of seizure onset.

**Conclusions:** Properties of interictal functional networks from scalp EEG can be estimated using a computer model and used to predict seizure likelihood and onset patterns. Our framework, consisting of three clinically-meaningful measures, could be implemented in the clinic to quantify the diagnosis and seizure onset pattern.

## Introduction

Brain function is increasingly understood in terms of large-scale brain networks. Disruptions to these networks can lead to a range of neurological conditions, including seizure disorders and epilepsy.^1^

In focal epilepsy, the conventional concept is that a single abnormal brain region generates seizures. However, this concept does not explain the re-emergence of seizures after apparently successful surgical resection of the presumed seizure focus,^2^ or that failure of epilepsy surgery can be predicted by features of an extended brain network beyond the seizure focus.^3–7^ In generalised epilepsy, the conventional concept is that seizures emerge without focal onset. However, this concept does not explain focal driving nodes in animal models of generalised spike-wave discharges.^8^ We previously proposed a theoretical framework that could reconcile these observations, showing the interplay between dynamics in localised brain regions and the pattern of connections between them is fundamental to whether a brain network can generate seizures, and whether seizures appear focal or generalised.^9^

Recognising that the historic dichotomy or either focal or generalised onsets does not reflect the richness of epilepsies presenting clinically, classification schemes have been evolving to reflect this new understanding of large-scale network mechanisms.^10–11^ A rigorous, quantitative, framework is required to underpin these new classifications of focal, generalised, combined focal and generalised and unknown epilepsies, in particular to quantify: (i) the propensity of a brain to generate seizures; (ii) how localised is the generation of a seizure; and (iii) how seizure activity propagates through large-scale brain networks.

In the epilepsy clinic, EEG is ubiquitous and provides a means to observe large-scale brain networks, and observing seizures and interictal discharges in EEG underpins the diagnosis and classification of epilepsy. Under the assumption that the properties of large-scale brain networks are a critical component of the onset of seizures and interictal discharges, alterations in network properties should be an enduring feature of the seizure prone brain and thus reflected within EEG regardless of whether seizures or interictal discharges are present. In support of this concept, we have developed methods for extracting functional network properties from 20s segments of resting-state EEG,^12,13^ and shown how properties of these networks are altered in people with idiopathic generalised epilepsy (IGE).^14^

Here we provide an objective method to quantify large-scale brain network features, using short epochs of apparently normal resting-state EEG. We introduce the onset index (OI), as a measure of the ability of a brain region to drive seizure onset, and the participation index (PI), as a measure of the ability of a brain region to become involved in ongoing seizures. We use these measures to test whether the pattern of interictal brain network connections is a major determinant of the capacity of a brain to generate seizures, and quantify the degree to which the pattern of seizure dynamics appears focal or generalised.

## Methods

### Subject Recruitment and Data Collection

We recruited 38 healthy subjects with no history of neurological or psychiatric disorders, and 68 subjects who had an established diagnosis of epilepsy (in accordance with International League Against Epilepsy guidelines and criteria): 25 with idiopathic generalised epilepsy, 23 with left focal epilepsy, and 20 with right focal epilepsy. People with epilepsy were recruited between March 2011 and October 2014 from the epilepsy clinics at King's College Hospital, London UK, and were a consecutive series who fitted the inclusion and exclusion criteria and were able to participate. Adult patients over 18 years of age were recruited, with epilepsy currently treated with AEDs, ongoing seizures (at least one per year), and no other neurological or psychiatric disorders (see Supplementary Materials). Healthy subjects were recruited from a local volunteer database. The outcomes in this study were a functional network derived from the interictal EEG of each subject, and a model-derived prediction of the pattern of seizure onset based on this network. We compared the model predictions with clinical classification of seizure onset (focal or generalised).

All EEG recordings were collected in the Department of Clinical Neurophysiology at King's College Hospital. Ag-AgCl (10 mm) disc electrodes were fixed at scalp positions in the modified Maudsley configuration.^15^ Ground and reference electrodes were placed between Pz and Cz and Cz and Fz, respectively. EEGs were recorded using Nicolet amplifiers (Viasys Healthcare, San Diego, California, USA). Data were collected at a sampling rate of 256Hz with filters set at 0.5 and 70Hz. Impedances did not exceed 5kOhms.

For this study one 20-second segment of eyes closed, resting-state EEG (subjects awake, eyes closed, no interictal discharges or seizures, minimal artefacts) was extracted for each subject by an EEG trained clinician (FAC and SJ). The epoch duration of 20 seconds is a compromise between availability of artefact-free segments in routinely collected EEG in clinics, and robustness of functional network properties.^16,17^ The interictal EEG segments were band-stop filtered between 48-52Hz to exclude power line interferences. We then computed the average power across all channels and used this value to normalise the time series from each channel, rather than normalising each channel individually. This has the advantage that relative differences in power between channel locations are preserved. Finally, we used a bandpass filter to extract data in the low alpha (6-9Hz) frequency band.^18^ This choice of frequency band has been demonstrated previously to be at the basis of significant differences between generalised epilepsies and healthy controls in resting-state EEG.^12-14,19^ We emphasise here that our analysis of EEG was based entirely on apparently normal, resting-state EEG free from interictal discharges, seizures or artefacts. This is a critical point. Seizures emerge from non-seizure brain states, therefore understanding the ability of non-seizure brain states to support transitions to seizure is crucial.

### Extraction of Functional Networks from EEG

Following (Schmidt *et al*., 2014), we derived a functional network for each subject from their segment of resting state EEG, using a correlation-based method called the Phase Locking Factor (PLF).^20^

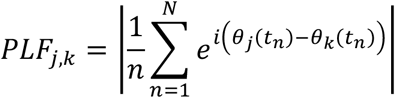

The PLF is a measure of the level of synchrony between two EEG channels and was computed by finding the maximum level of pairwise correlation between the instantaneous phases of these two channels (*θ*_*j*_, *θ*_*k*_). By applying this method to all possible combinations of channel pairs, we obtained a weighted functional network consisting of 19 nodes and connectivity strengths between the nodes defined by the corresponding values of the PLF. Contributions at zero phase-lag were rejected in order to minimise the problem of common sources and volume conduction.^21^

In order to detect and reject spurious connections between nodes (i.e. significantly different than noisy uncorrelated time-series), we additionally carried out a surrogate-analysis. We created 99 surrogate networks for each segment of resting-state EEG, using a Fourier-based method which preserved the autocorrelation and spectrum of the original time.^22^ A connection from the original functional network was rejected (i.e. the connectivity strength set to 0) if the corresponding connection was found to be significantly stronger in the surrogate networks.

### Dynamic Network Model of Seizure-like Activity in the Brain

With increasing interest in functional networks over recent years, a new approach has emerged – which we term a ‘dynamic network model’ – that places a mathematical description of critical features of brain dynamics (e.g. seizures or the transition to seizure) on each node, with connection strengths between nodes governed by an overall functional network. Herein, a Kuramoto model,^23^ which describes the dynamics of a single population of N identically coupled phase oscillators, was used for each node. Phase oscillator models have been increasingly used in a neuroscience setting.^12,24–26^ We extend the standard model for a single population of N phase oscillators, to a network consisting of M populations:

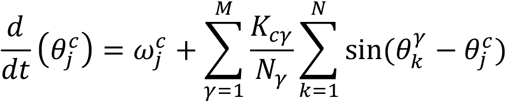

with phase *θ*, natural frequency 𝜔, and coupling strengths K_c_γ. Here the number of populations M is equal to the number of ‘sensors’ (i.e. M=19 (EEG channels)) and the number of phase oscillators N in population *γ* is assumed to be very large, since a single EEG channel measures the collective electrical activity of a large neuronal population.

By carrying out a standard transformation, the order parameter r_c_ of a population of Kuramoto oscillators can be calculated. r_c_ takes values between 0 and 1, where 0 represents an asynchronous, low-amplitude (non-seizure) state and 1 corresponds to a fully synchronised, high-amplitude (seizure-like) state throughout the population. For r_c_ between 0 and 1, the levels of coherence and amplitude increase. Finally, by averaging across all the values of r, we obtain a global order network parameter r_g_, which quantifies the amount of synchronisation over the network of M populations. Stationary solutions for these equations can be computed numerically and analytically.^12^

### Global Mechanism of Seizure Onset

To characterise the contribution of network connectivity to the ability of a network to generate seizures, we introduce an additional global coupling parameter C which scales the adjacency matrix of PLF values:

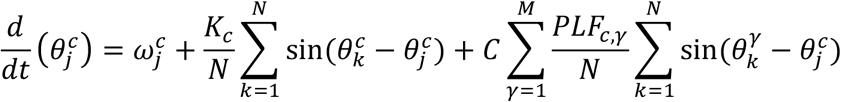

Note there are two types of coupling in this model: K_c_ governs the coupling within a separate population c, whereas the PLF-values determine the coupling between populations. As C increases, individual regions within the network may make the transition to their synchronous state. We term the value of C for which the network transitions to its synchronised state the critical coupling value (C_c_). In previous work, we found this critical coupling value to be significantly lower in a cohort of subjects with IGE in comparison to healthy controls.^12,13^ This indicates that the resting state functional networks of people with IGE support transitions to seizures more readily than those from healthy controls. Here, we can regard the critical coupling value as a generic marker of the propensity of a brain to generate seizures of any type (focal or generalised). We calculated the critical coupling value C_c_ for every subject and performed pairwise comparisons at the group-level.

### Local Mechanisms of Seizure Onset and Participation

To characterise the contribution of individual brain regions to whether seizure patterns appear focal or generalised, we introduced two new measures for characterising dynamic network models: the onset index (OI) and the participation index (PI). Both indices can be calculated directly from the ‘activation matrix *ACT*’ with entries *ACT*_*j,k*_ ∈ [0,1], quantifying the amount node j spent in the seizure-like state a result of node k being in the seizure-like state (see Figure 2). Mathematically we have: ACT_j,k_ = T_j_/T_total,k_, with T_j_ the total time node j spent in the seizure-like state and T_total,k_ the total time of simulation where node k was in the seizure-like state at t=0. In particular, *ACT*_*j,k*_ = 0 means that node j was not driven into the seizure-like state by the seizure-like activity in node k, whereas node j spent the entire simulation in the seizure-like state if *ACT*_*j,k*_ = 1.

**Figure 1:**
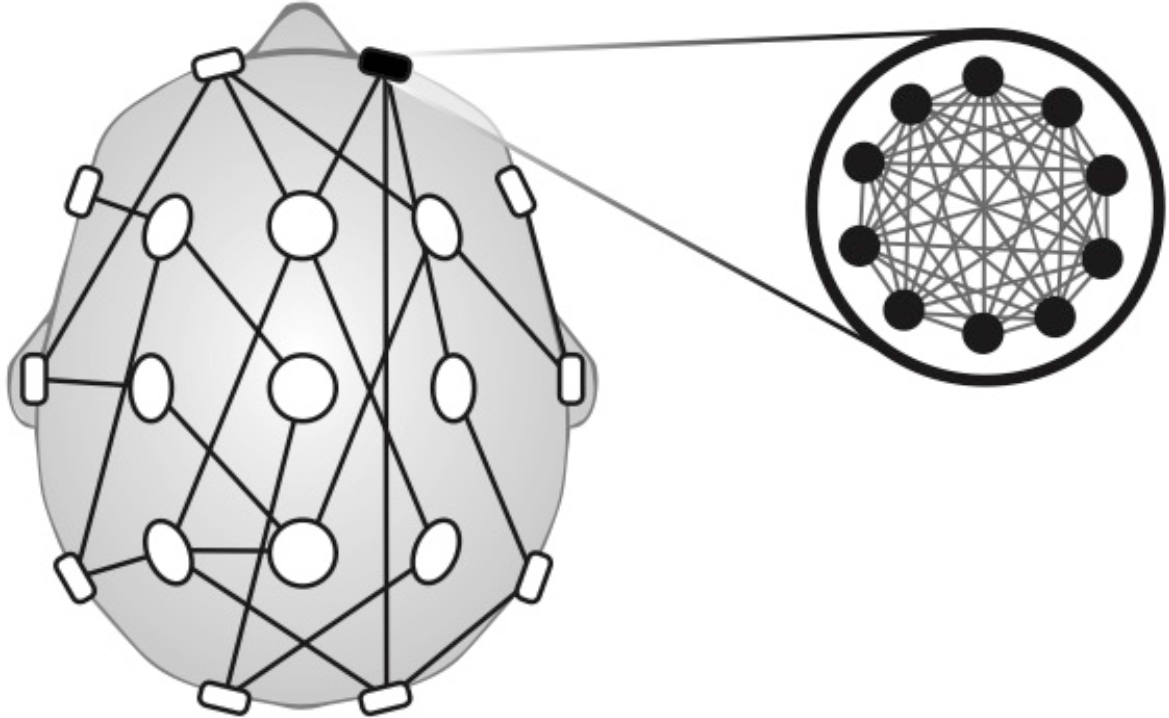
Schematic of Dynamic Network Model. Schematic drawing of a particular dynamic network model: each individual region is a model representation of coupled Kuramoto oscillators, and the connectivity between the regions is determined by the PLF values.

**Figure 2:**
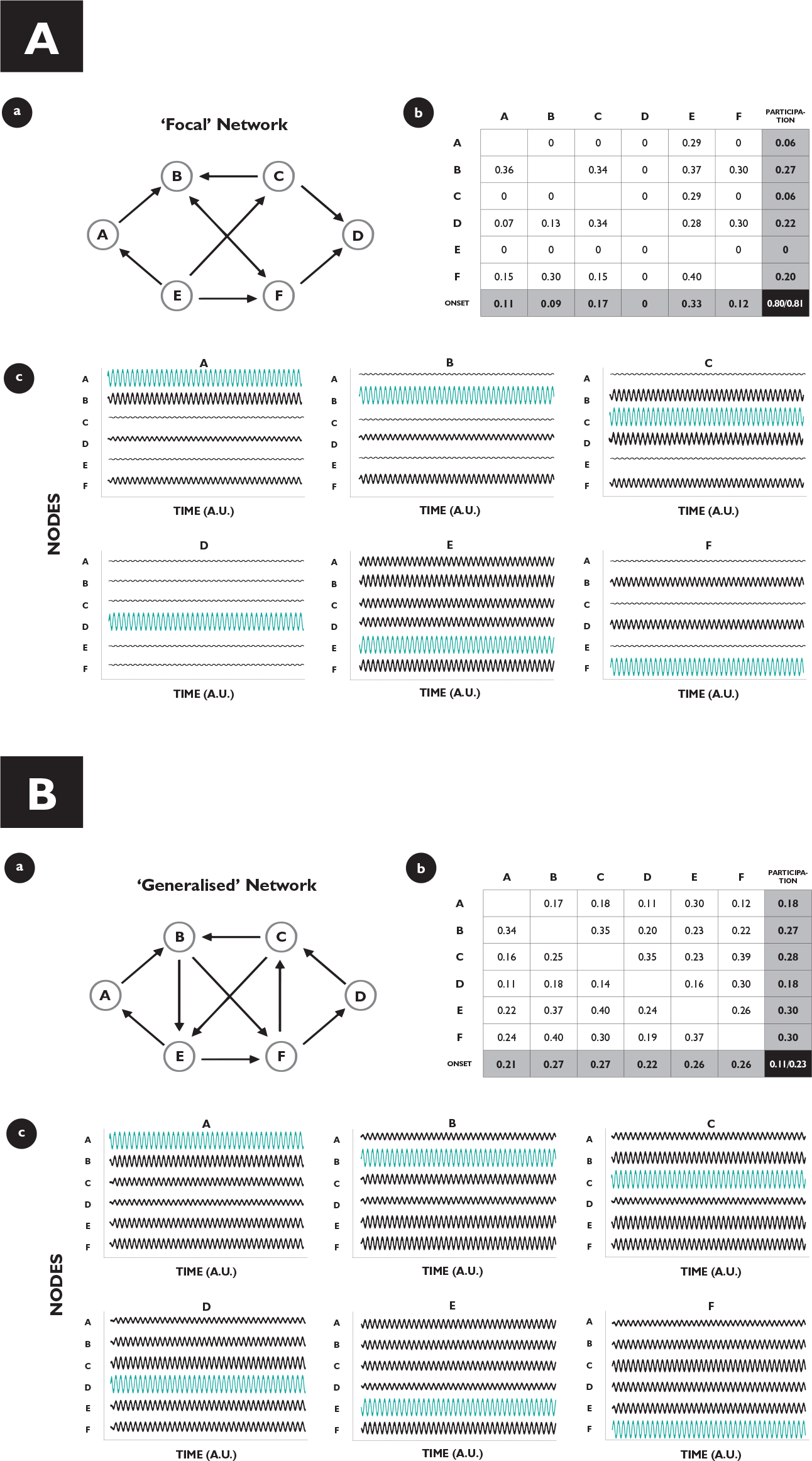
Characterising the Focal or Generalised Nature of a Brain Network. **A)** Onset index (OI) and participation index (PI) for a “focal” network. a) Network structure with 6 nodes (A, B, C, D, E, F) and edges describing directed connectivity between the nodes. b) Activation Matrix for the given network. Each node is set into the locally synchronised state once, and its response of the remaining nodes calculated (which constitute the entries of the activation matrix). The OI for a node I corresponds to the averaged column sum of column i, and the PI for a node i corresponds to the averaged row sum of row i. The variability in the OI is found by dividing the standard deviation over all the Onset Indices and dividing this by the mean over all the OI values of the network, and equivalently for the PI. The normalised standard deviation for the OI and the PI are shown in the bottom right of the activation matrix (light grey). c) The dynamics of each node corresponds to the collective activity of a subpopulation of Kuramoto oscillators and can be thought of as a single channel of simulated EEG, where low amplitude activity represents the non-synchronised state (interictal), and high amplitude oscillatory activity represents the synchronised state (ictal). In each subpanel, a node is set into the synchronised state (blue), and the network response simulated. Note that the response of the network is variable and depends on the location of seizure initiation, indicating a high degree of heterogeneity in the network’s behaviour. **B)** OI and PI for a “generalised” network. a) Network structure with 6 nodes (A, B, C, D, E, F) and edges describing directed connectivity between the nodes. b) Activation Matrix for the given network. Each node is set into the locally synchronised state once, and its response of the remaining nodes calculated (which constitute the entries of the activation matrix). The OI for a node I corresponds to the averaged column sum of column i, and the Participation Index for a node i corresponds to the averaged row sum of row i. The variability in the OI is found by dividing the standard deviation over all the OI values and dividing this by the mean over all the OI values of the network, and similarly for the PI. The normalised standard deviation for the OI and the PI are shown in the bottom right of the activation matrix (light grey). c) The dynamics of each node corresponds to the collective activity of a subpopulation of Kuramoto oscillators and can be thought of as a single channel of simulated EEG, where low amplitude activity represents the non-synchronised state (interictal), and high amplitude oscillatory activity represents the synchronised state (ictal). In each subpanel, a node is set into the synchronised state (blue), and the network response simulated. Note that the response of the network is homogeneous throughout, indicating a high degree of similarity across the network’s behaviour.

Averaging across the j-th column of the activation matrix, the OI of node j is obtained: a measure of the capacity of node j to drive synchronisation across the rest of the network. The OI is computed numerically by starting with a network where all nodes are in an asynchronous setting and then increasing the internal coupling strength of the j-th population of phase oscillators above a critical value such that the j-th population becomes synchronised and displays seizure-like activity. The response in the remaining nodes of the network to this localised seizure-like activity is then computed. The OI is the mean value of these responses and therefore takes a value between 0 and 1. Zero corresponds to a ‘disconnected’ node: even if it is in the seizure-like state, there is no node it was able to influence. In contrast: if a node has an OI equal 1, it recruited all the other nodes in the network into their seizure-like state. Here, we can regard the OI as a specific marker of the tendency of a brain region to generate seizures; we would expect that in focal epilepsy, OI would be variable between regions, reflecting that a particular localised subset of nodes drives seizure onset; whereas in generalised epilepsy we would expect OI to be less variable between regions, reflecting that in these cases a “seizure focus” is not expected.

Averaging along the j-th row of the activation matrix, we obtain the Participation Index (PI) of node j: a measure of the capacity of node j to be synchronised by other nodes in the network themselves being synchronous. The PI is computed numerically by calculating the amount of synchronisation in node j in response to increasing the internal coupling strength K of all other nodes *j* ≠ *k* above their critical value such that each population individually becomes synchronised. As for the OI, the mean value of the response to all other nodes is calculated and therefore the PI takes a value between 0 and 1. A node with PI equal 0 is classed as ‘disconnected’: none of the other nodes were able to recruit it into its seizure-like state. In contrast: a node with PI equal 1 is recruited into its seizure-like state by all other nodes within the network. Here, we can regard the PI as a specific marker of the tendency of a brain region to become involved in the seizure network; we would expect that in focal epilepsy, PI would be variable between regions, reflecting that a localised subset of nodes constitutes the seizure network; whereas in generalised epilepsy we would expect PI to be less variable between regions, reflecting that the seizure network is much more widespread.

### Identification of Side of Seizure Onset Using OI and PI

Since we can define the OI and the PI on a region-by-region basis within a functional network, we hypothesise that regions indicated by the OI and the PI as significantly different in the focal cohort, correlates with the clinically indicated hemisphere. In contrast, we hypothesise that within the IGE cohort there would be no regions statistically different from the control cohort, reflecting the apparent lack of specificity of onset and spread in generalised seizures.

### Statistical Analysis

Statistical analysis was carried out using MATLAB (version 17). Two nonparametric tests, the Kruskal-Wallis test and the Mann-Whitney U test (two-tailed, medians) were used throughout to determine statistical significance (p<0.05) between groups of an independent variable (critical coupling value, variance of OI, variance of PI). In particular, a Krusal-Wallis test was applied to test for statistical significance (p<0.05) between groups for an independent variable. If the Kruskal-Wallis test was found to be significant, then a Mann-Whitney U test was carried out to explicitly test between-group differences, and a Bonferroni correction was applied to correct for multiple comparisons (e.g. Control vs Focal, Focal vs IGE). All p-values computed for determining the side of seizure onset were corrected with a Bonferroni factor of 57: since three group comparisons were carried out against healthy controls (Left Focal, Right Focal, IGE) for every brain region (19 in total).

### Patient Consent

The study was approved by Bromley Research Ethics Committee (reference 12/LO/0230) and all subjects gave their informed written consent to take part

## Results

We analysed 20s epochs of resting-state EEG collected from 106 subjects: 43 subjects with focal epilepsy (23 left focal, 20 right focal), 25 subjects with idiopathic generalised epilepsy, and 38 healthy controls.

### Critical Coupling Value

We found that the effect of progressively increasing the global coupling strength between brain regions ultimately led to widespread synchronised activity. We term the value for which the first brain region becomes synchronised the critical coupling value. Using a Kruskal-Wallis test (p<0.05), we found that the critical coupling value differed between groups (Figure 3, Table 1). We then examined pairs of groups using a MWU-test (medians, two-tailed, p<0.05, Bonferroni correction: x3): the dynamic network model when the network was inferred from people with epilepsy was more prone to generate seizure-like activity than those from healthy controls: that is, the critical coupling values from people with epilepsy were found to be significantly lower in contrast to the values from the healthy control cohort. We found no significant difference for the mean critical coupling value between the focal cohort and the generalised cohort.

**Table 1:** Critical Coupling

Kruskal Wallis: $p=0.00002$
| **Group-comparison** | **p-value (MWU)** | **Bonferroni corrected (x3)** |
| --- | --- | --- |
| Con – IGE | 0.0014 (U = 703) | 0.0041 |
| Con – Focal | <0.0001 (U = 1261) | <0.0001 |
| Focal – IGE | 0.0781 (U = 399) | 0.2344 |

**Figure 3:**
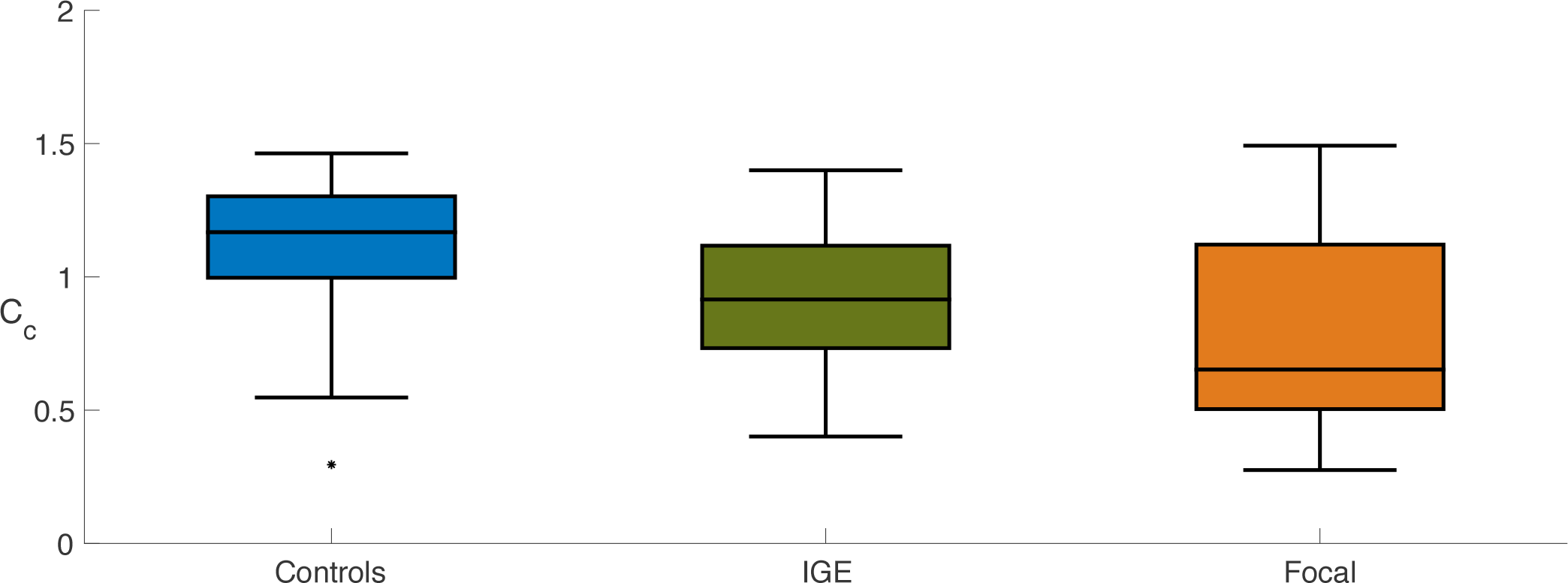
Group Comparison for Critical Coupling Value. Boxplots displaying the distribution of the critical coupling values for the control subjects (n = 38, median = 1.1674, IQR = 0.3055), generalised epilepsies (n = 25, median = 0.9152, IQR = 0.3843) and focal epilepsies (n = 43, median = 0.6520, IQR = 0.6170): the box is spanned by the first quartile to the third quartile (interquartile range IQR), with the median (bold line) situated within. Whiskers above and below the box show the location of the largest and smallest value within 1.5 times the IQR, and suspected outliers are depicted as stars. Mann-Whitney U tests rejects the null-hypothesis that the critical coupling for the focal and generalised epilepsies have the same distribution as the controls (Controls – Generalised: p = 0.0041, U = 703; Controls – Focal: p < 0.0001, U = 1261; medians, two-tailed, p<0.05, Bonferroni correction: x3).

### Onset Index

The OI measures this ability of a brain region to recruit the rest of the network into the seizure-like state, and displayed on average higher levels of variance across all brain regions in the focal cohort in comparison to the generalised cohort. In particular, we found that the response of an individual brain to the onset of synchronised activity within a localised brain region was statistically significantly different between generalised and focal cohorts (p = 0.0056; U = 319; Figure 4a; MWU-Test, medians, two-tailed, p<0.05). In networks from the generalised cohort, most brain regions could drive the onset of seizure-like behaviour. In contrast, we found that in networks from the focal cohort there was typically only a small number of brain regions that could drive the onset of seizure-like behaviour. For these networks, the onset of large-amplitude, synchronised activity in most regions would not significantly affect the activity of the rest of the network.

**Figure 4:**
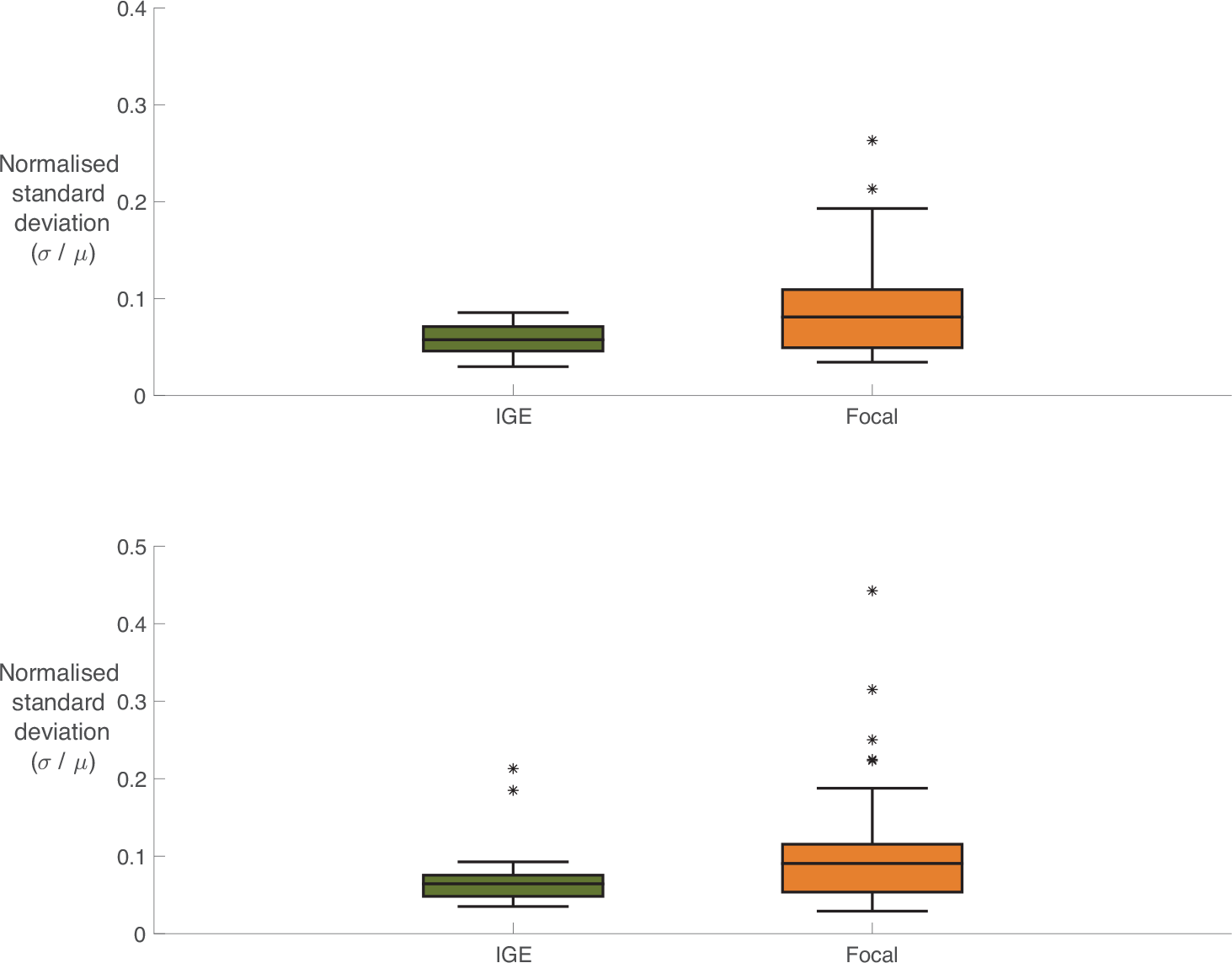
Comparing Variability in OI and PI for Focal and Generalised. **A)** Boxplots displaying the distribution of normalised standard deviation of the OI comparing focal epilepsies (n = 43, median = 0.0808, IQR =0.06) against generalised epilepsies (n = 25, median = 0.0573, IQR = 0.0252). Specifically, the normalised standard deviation is calculated by dividing the standard deviation of the OI across all the nodes within the network by the mean of the OI across all the nodes within the network: 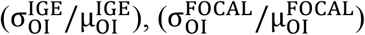. Mann-Whitney U test rejects the null-hypothesis that the normalised standard deviation of the OI for the focal and generalised epilepsies have the same distribution (p = 0.0056, U = 319, medians, two-tailed, p<0.05). **B)** Boxplots displaying the distribution of normalised standard deviation of the Participation Index comparing focal epilepsies (n = 43, median = 0.0905, IQR = 0.0618) against generalised epilepsies (n = 25, median = 0.0642, IQR = 0.0273). The normalised standard deviation is calculated by dividing the standard deviation of the Participation Index across the nodes within a network by the mean of the Participation Index across the nodes within that network: 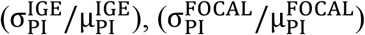. Mann-Whitney U test rejects the null-hypothesis that the normalised standard deviation of the PI for the focal and generalised epilepsies have the same distribution (p = 0.0206, U = 355, medians, two-tailed, p<0.05).

### Participation Index

The PI measures the response of a brain region to seizure-like activity initiated from another region within the network and showed higher levels of standard deviation in the focal cohort in comparison to the generalised cohort. In particular, we found that the response of other brain regions to the onset of abnormally synchronised activity within a localised brain region was different in generalised and focal networks (p = 0.0206; U = 355; Figure 4b, Mann Whitney U Test, medians, two-tailed, p<0.05). In the generalised cohort, most brain regions became involved in ongoing seizure-like activity instigated at some other place within the network, regardless of the location of the localised brain region. In contrast, we found that for focal networks the response of other brain regions to the onset of seizure-like activity was heterogeneous. Typically, seizure-like activity remained confined to a smaller cluster or a subnetwork of the larger global brain network.

### Side of Seizure Onset

Given that the distribution of the OI in the focal epilepsy cohort was non-uniform, we explored the specific values of the OI on a region-by-region basis, hypothesising that the hemisphere with highest OI would be concordant with the side of the clinically determined seizure onset. In particular, we found specific brain regions within the focal networks that were statistically significant drivers of seizure-like behaviour in comparison to the control cohort. In the cohort with left focal epilepsies, we found several regions in both hemispheres that displayed a significant difference in the OI (Figure 5, Table 2) in contrast to the control cohort. However, the region with the strongest level of significance was found in the left hemisphere. In the cohort with right focal epilepsies, we found there was one region in the right hemisphere that displayed a significant increase in the OI in contrast to the control cohort. For the cohort of generalised epilepsies, no such differences were found between OI in any region and OI in the same region in the control cohort. This was consistent with our previous observation that most regions could drive the onset of seizure-like behaviour in networks from the generalised cohort. A similar analysis for the PI revealed that there were several significantly different regions for the left focal epilepsies, and the region that was most strongly different was situated on the left side. Similarly, for the subjects with right focal epilepsies, we found one region in the right hemisphere that displayed a significant increase in contrast to the control cohort. For the cohort of generalised epilepsies, no differences were found for any region, which is consistent with our previous observation that a large set of regions becomes involved in seizure-like behaviour in networks from the generalised cohort.

**Table 2:** Significantly Different Regions in Left Focal, Right Focal, and IGE Left Focal

Left Focal
| **Onset Index** |  | **Participation Index** |  |
| --- | --- | --- | --- |
| **Channel** | **p-value (BC: 19x3)** | **Channel** | **p-value (BC: 19x3)** |
| F7 | 0.030 (U = 204) | Fp2 | 0.0181 (U = 195) |
| C3 | 0.035 (U = 207) | F3 | 0.0202 (U = 197) |
| T3 | 0.049 (U = 213) | T5 | 0.0128 (U = 189) |

Right Focal
| **Onset Index** |  | **Participation Index** |  |
| --- | --- | --- | --- |
| **Channel** | **p-value (BC: 19x3)** | **Channel** | **p-value (BC: 19x3)** |
| P4 | 0.048 (U = 176) | Fp2 | 0.0182 (U = 160) |

**Figure 5:**
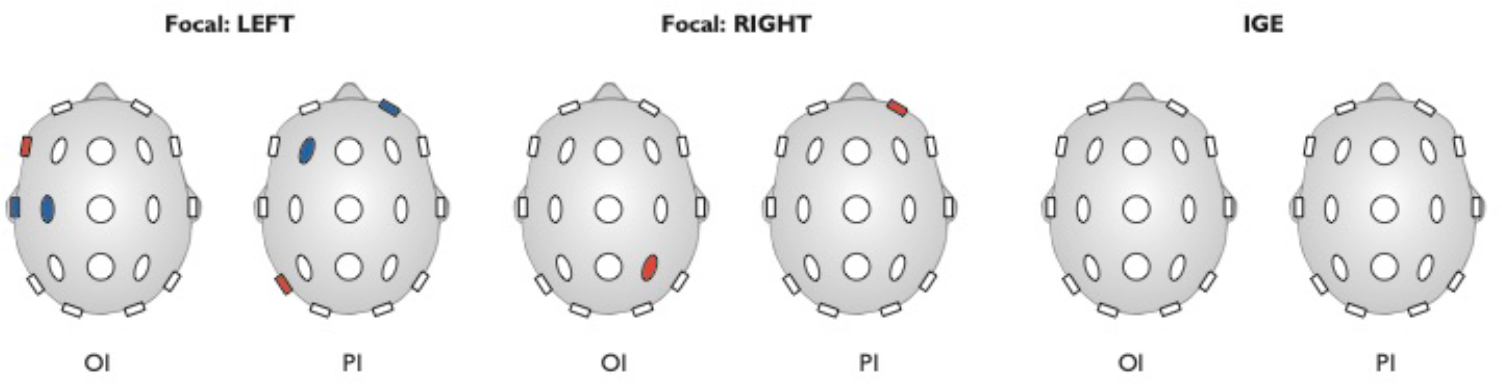
Determining Side of Seizure Unset Using OI and PI. Regions in blue indicate a significant (Mann-Whitney U Test, two-tailed, p<0.05) difference between the distributions of normalised standard deviations of a group (Left Focal (n = 23), Right Focal (n = 20) or IGE (n = 25)) and the control cohort (n = 38) for that specific region. Regions in red indicate the strongest significant difference of all regions in the network. For the regions in black the outcome of the Mann-Whitney U test was to accept the hypothesis that the distributions are similar for that region. We found no regions for the generalised epilepsies that were significantly different to the healthy controls, whereas for focal epilepsies the regions with highest values of normalised standard deviations of the OI and PI were associated with the affected hemisphere.

## Discussion

The concepts of focal and generalised epilepsy have changed considerably in recent times, moving towards a conceptual framework based on the manner in which seizures emerge in and engage localised versus widely distributed brain networks. At the current time, these network concepts remain qualitative and lack a robust descriptive, quantitative framework. Here, we provide an approach that allows network origins of seizures to be described in terms of three parameters: global critical coupling, which describes the propensity of a brain to generate any seizures; onset index (OI), which described the tendency of seizures to generate from specific regions; and Participation Index (PI), which describes the tendency of seizures to engage specific networks. This simple, intuitive framework provides a robust and highly flexible way to precisely define the meaning of “focal” and “generalised” in any specific example of epilepsy. We believe this approach reveals fundamental mechanistic phenomena of epilepsy and provides a future tool for clinical classification of seizures and epilepsy.

Here, we applied this framework to the most readily-available diagnostic modality in epilepsy – conventional 19 channel scalp EEG. A computational dynamic network model with parameters inferred from a 20s segment of interictal, resting-state EEG enabled the characteristic brain network properties of healthy control, generalised, and focal subjects to be described. First, we found that resting-state brain networks of people with either focal or generalised epilepsy are situated closer to a transition between normal activity and seizure-like activity. This was shown by observing, in both the focal and generalised cohorts, lower values of the critical coupling - that is the coupling strength of the global network for which an individual brain region is driven into a seizure-like state. To understand mechanistically why the pattern of electrical activity at the level of the EEG appears focal or generalised, we introduced two new measures - the onset index (OI) and participation index (PI). The OI characterises the ability of a given brain region to drive a seizure within an overall brain network. We found that the OI is less uniformly distributed across regions within brain networks from the focal epilepsy subjects in comparison to subjects with generalised epilepsy. This confirms a tendency for the onset of seizure-like activity from resting-state brain networks to be localised in the case of focal epilepsy but not in the case of generalised epilepsy. Furthermore, when the focal cohort was considered as subgroups of left focal epilepsies and right focal epilepsies, higher variability of OI associated with the affected hemisphere. The PI characterises the tendency for a given brain region to become involved in a seizure driven from another brain region. As with OI, we found that the PI is less uniformly distributed across regions within brain networks from the focal epilepsy subjects in comparison to generalised epilepsy subjects. When the focal cohort was considered as subgroups of left focal epilepsies and right focal epilepsies, higher variability in PI associated with the affected hemisphere. Together these results reveal from background activity in scalp EEG that focal seizures preferentially engage specific localised networks rather than the entire brain and suggests a tendency for seizures in generalised epilepsy to engage widely distributed brain networks, in keeping with current concepts and evidence.^10,27,28^ In summary, our findings suggest that brain networks supporting generalised seizures are more homogeneous with similar driving tendency and network engagement responses across most regions within a network, whereas networks supporting focal seizure contain heterogeneity and are typically asymmetric.

From a diagnostic perspective, the presence of epileptiform abnormalities in EEG remains currently the most useful biomarker of epilepsy. However, even during long-term video-EEG an individual with epilepsy may not display epileptiform abnormalities, giving rise to a well-known problem of false-negatives in diagnostic EEG, as well as a low incidence of false positives in healthy subjects.^29^ Additionally, over-reading of EEG is a further cause of misdiagnosis.^30^ Misdiagnosis and mistreatment of epilepsy is a serious problem with significant negative consequences for the subjects involved and carries a significant financial burden.^31,32^ Consequently, a data-derived biomarker from interictal EEG recordings offers the potential to significantly support current clinical practice by providing a quantitative framework for diagnosis of both focal and generalised epilepsies.

A key feature of our study is the ability to reveal network markers of seizures from short epochs of interictal, resting-state, 19 channel scalp EEG, which is very commonly available in epilepsy centres worldwide. We used an established data-driven modelling approach with minimal assumptions about the underlying properties of the recorded signals (e.g. stationarity) in order to provide a quantitative account of focal and generalised epilepsy. Finding evidence for focal and generalised network features in resting-state interictal EEG suggests that the causal network properties that drive seizure onset and propagation are observable even in the absence of seizures and interictal discharges; in other words, in epilepsy the brain has an enduring trait to produce seizures of specific types.^33^ Furthermore, and importantly, revealing these features in a short segment of normal resting-state interictal 19 channel scalp EEG opens a novel opportunity to diagnose and classify epilepsies without observing seizures; we do not claim here to have proven the clinical value of this method, but have provided a key foundation for future work.

At the same time, our use of 19 channel scalp EEG imposes limitations. Our approach is intrinsically confined to a ‘sensor space’ analysis and consequently we cannot infer any causal relations between the processes underlying the recordings. In line with this, it is important to note that the computer model used is an abstract, phenomenological description of the recorded EEG signals that has no direct neurobiological interpretation. Other modalities such as MEG and fMRI have been used to study the network mechanisms underlying seizure generation, in particular focal seizures.^34,35^ In the future, high-density EEG may become more widespread in clinical settings which would justify more detailed approaches such as source-based reconstruction with a neurophysiologically detailed computer model. Such approaches hold further potential in providing support in diagnosing and lateralising epilepsy outside of the standard, clinical environment.^36^ Here, we have not attempted to take into account whether the subjects with focal epilepsy had secondarily generalization or not; a new and larger set of data would allow us to examine whether these methods are sensitive to differentiating subjects whose focal seizures generalize secondarily very rapidly from those for who this process occurs slower. Additionally, useable data for our study was limited by the fact that an EEG-trained clinician was required to select the EEG epochs for further analysis. This might be automated in the future, but in our current analysis could be considered to provide a risk to the robustness of the presented measures. A further, potential, confounding influence is the effect of time of day on resting-state EEG. Factors such as circadian rhythms and environmental changes are known to dynamically influence cortical excitability.^37^ Finally, since there is no evidence that one of the patient groups (focal or generalised) is significantly more affected by AEDs or drug-load, it is unlikely that this could lead to an unbalanced effect on the outcome measures we present.

Within a brief epoch of normal background EEG we can find imprints of the fundamental properties related to the overall susceptibility for seizure occurrence. In particular, these properties are fundamentally different between focal or generalised seizures, which are characterised objectively using only three features of a simple mathematical model.

## Contributors

WW designed the study, did the literature search, developed the model, analysed the data, made the figures and wrote the report. HS designed the study, developed the model, analysed the data, made the figures and wrote the report. EA, FAC, ADP and SJ recruited patients and acquired data. MPR designed the study, did the literature review and wrote the report. JRT designed the study, developed the model and wrote the report. All authors approved the final content of the report.

## Acknowledgements

We thank Lina Nashef PhD and Robert D.C. Elwes PhD for assisting in patient recruitment. We thank Marinho A. Lopes PhD for useful comments on the manuscript.

## Supplementary materials: Characteristics of the study participants

Clinical and Demographic Characteristics of Study Participants.

Clinical and Demographic Characteristics of Study Participants
|  | Subjects | Female | Age in years<br>(SD)*; range | Duration of<br>epilepsy in<br>years (SD);<br>range | Number of<br>AEDs (n) |
| --- | --- | --- | --- | --- | --- |
| **Controls** | 38 | 19<br>(50%) | 29.9 (8.6);<br>18-52 | - | - |
| **IGE** | 25 | 15<br>(60%) | 32.5 (11.1);<br>20-61 | 19.5 (14.7);<br>1-58 | 1 (13), 2 (9),<br>3 (3) |
| **Focal** |  |  |  |  |  |
| **- Left** | 23 | 9<br>(39%) | 38.7 (16.5);<br>20-77 | 15.4 (14.4);<br>2-55 | 1 (9), 2 (11),<br>3 (3) |
| **- Right** | 20 | 13<br>(65%) | 35.9 (13.2);<br>18-68 | 20.4 (11.2);<br>2-40 | 1 (6), 2 (10),<br>3 (4) |

*Caption: Data are presented as integers n, integers n (%), or mean (SD). The Mann-Whitney U test was used to compare age, duration of epilepsy and drug load between the groups (all tests failed to reject the null-hypothesis of equal medians of the groups, two-tailed, $p < 0.05$ ); Fisher's exact test was used for comparing gender (all tests failed to reject the null-hypothesis, $p < 0.05$ ).*

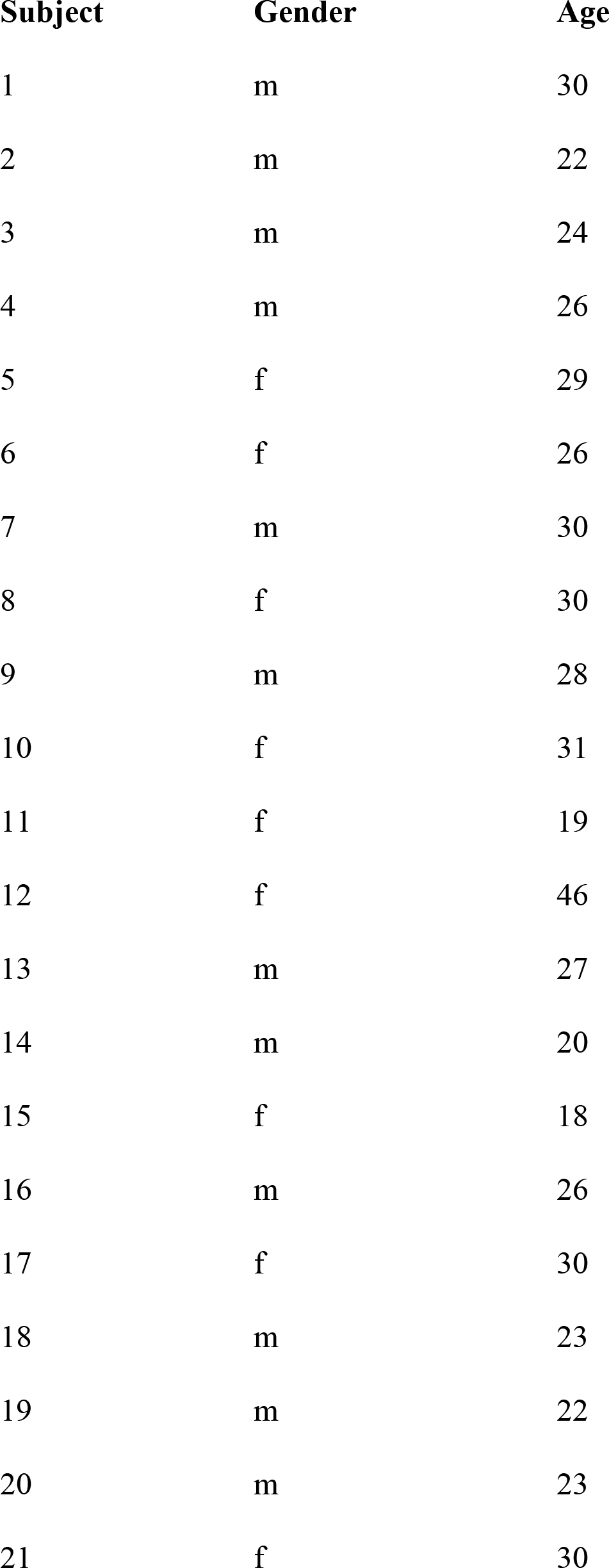
Control cohort.

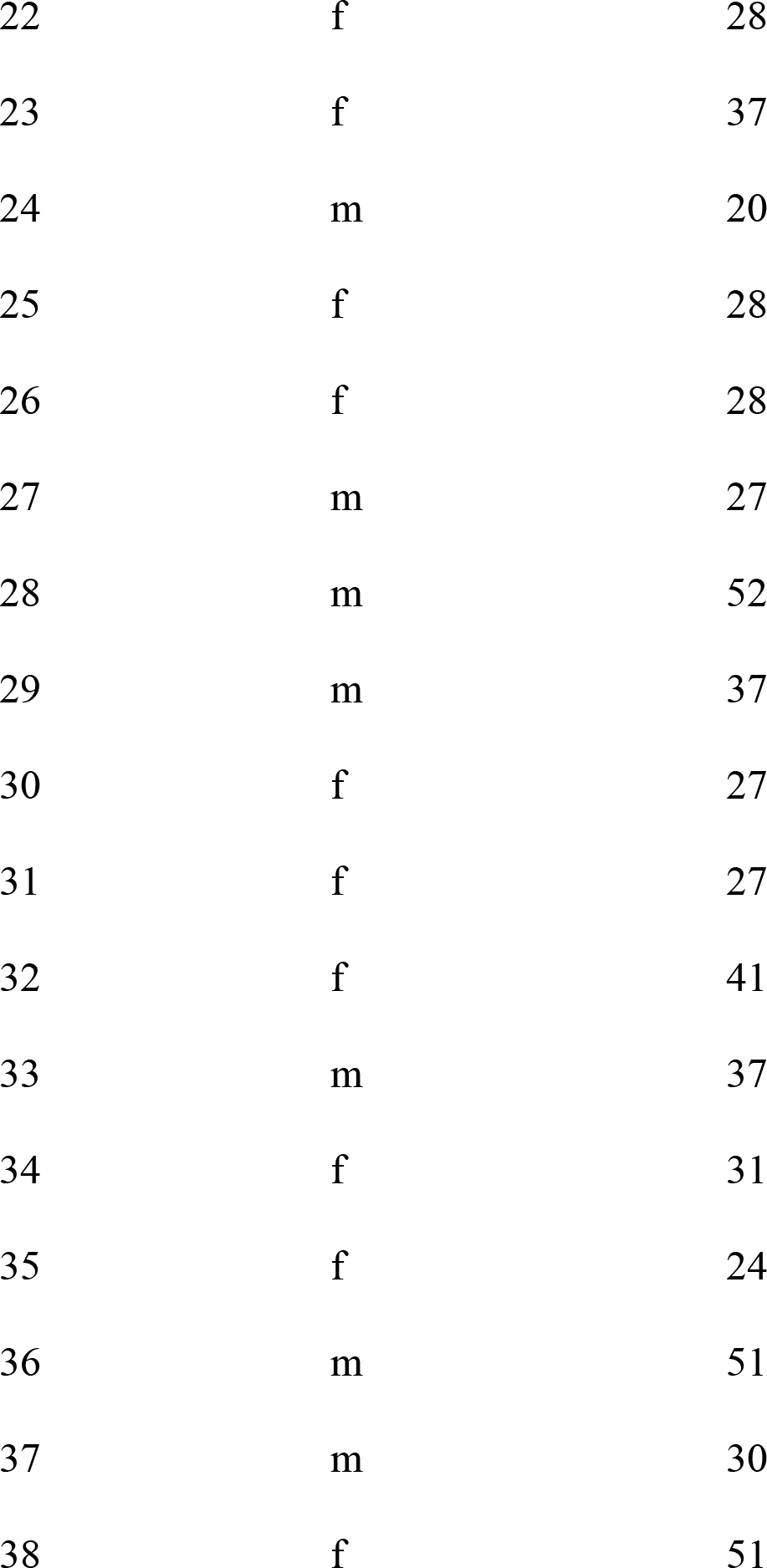

Idiopathic Generalised Epilepsy Cohort.

Idiopathic Generalised Epilepsy Cohort
| Subject | Gender | Age | Duration | Syndrome | Type of AED | MRI | EEG: | EEG: | EEG: | EEG: |
| --- | --- | --- | --- | --- | --- | --- | --- | --- | --- | --- |
|  |  | (years) | of disease<br>(years) |  |  |  | Background | Focal | focal | gen. |
|  |  |  |  |  |  |  | Slowing | Slowing | IEDs | IEDs |
| 1 | m | 31 | 17 | GTCS | CBZ | Normal | - | - | - | - |
| 2 | f | 40 | 31 | JAE | LTG, VPL | Normal | - | - | - | - |
| 3 | f | 31 | 5 | IGE | LTG, LEV | Normal | - | - | - | - |
| 4 | f | 52 | 50 | IGE | VPL, TPM | Normal | - | - | Yes | - |
| 5 | f | 47 | 36 | JAE | LEV | Normal | - | - | - | Yes |
| 6 | m | 25 | 21 | IGE | VPL | Normal | - | - | - | Yes |
| 7 | m | 25 | 14 | IGE | VPL, TPM | Normal | - | - | - | Yes |
| 8 | f | 28 | 8 | GTCS | CBZ | Normal | - | - | - | - |
| 9 | f | 21 | 11 | JAE | LTG, ETX | Normal | - | - | - | - |
| 10 | f | 39 | 31 | IGE | LEV | Normal | - | Yes | - | - |
| 11 | f | 21 | 15 | IGE | LTG | Normal | - | - | Yes | Yes |
| 12 | f | 20 | 5 | JME | LTG, LEV | Normal | - | - | Yes | Yes |
| 13 | m | 25 | 9 | IGE | VPL | Normal | - | - | - | - |
| 14 | m | 50 | 42 | CAE | VPL, LEV | Normal | - | - | - | - |
| 15 | m | 45 | 24 | CAE | VPL, LEV, LTG | Normal | - | - | - | Yes |
| 16 | m | 28 | 20 | IGE | VPL | Normal | - | - | - | Yes |
| 17 | f | 28 | 12 | IGE | LTG | Normal | - | - | Yes | - |
| 18 | f | 33 | 18 | IGE | VPL | Not available | - | - | - | - |
| 19 | f | 28 | 12 | IGE | TPM | Not available | - | - | - | Yes |
| 20 | m | 27 | 1 | IGE | VPL | Normal | - | - | - | Yes |
| 21 | m | 27 | 10 | IGE | CBZ, LEV, ZNS | Normal | - | - | - | Yes |
| 22 | f | 23 | 10 | IGE | VPL, LTG | Normal | - | - | - | Yes |
| 23 | m | 20 | 3 | IGE | TPM | Normal | - | - | - | Yes |
| 24 | f | 38 | 24 | IGE | LTG, LEV | Not available | - | - | Yes | Yes |
| 25 | f | 61 | 58 | IGE | LEV, CLB | Not available | Yes | - | Yes | Yes |

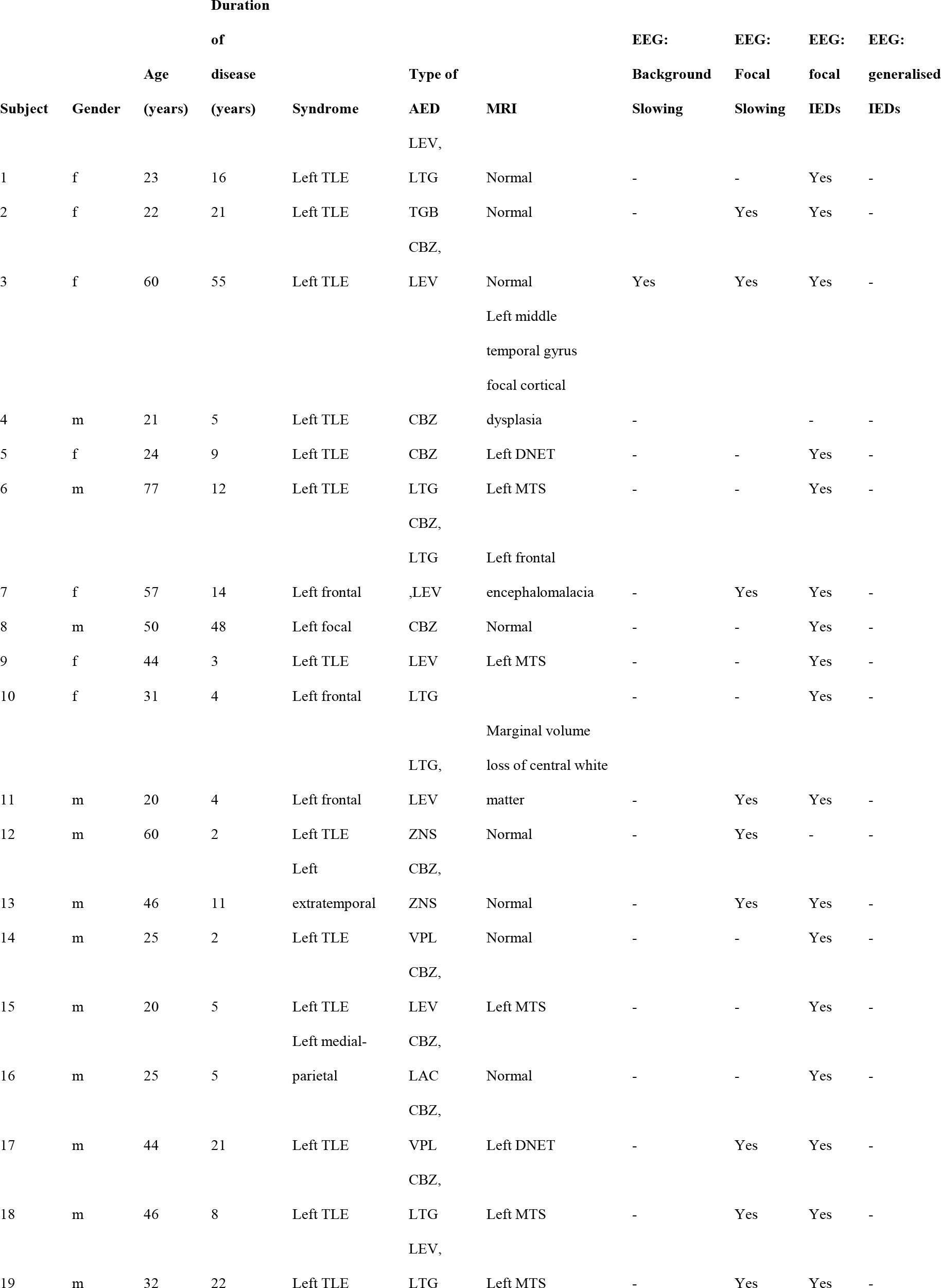
Left Focal Epilepsy Cohort.

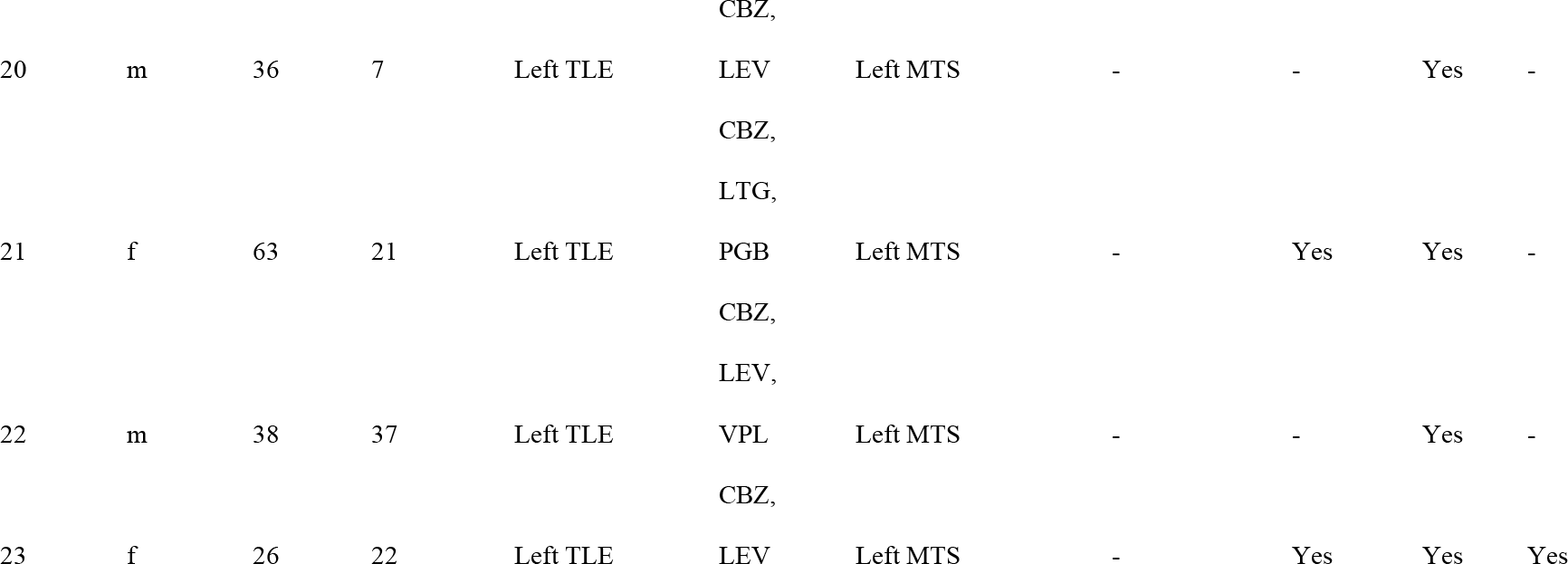

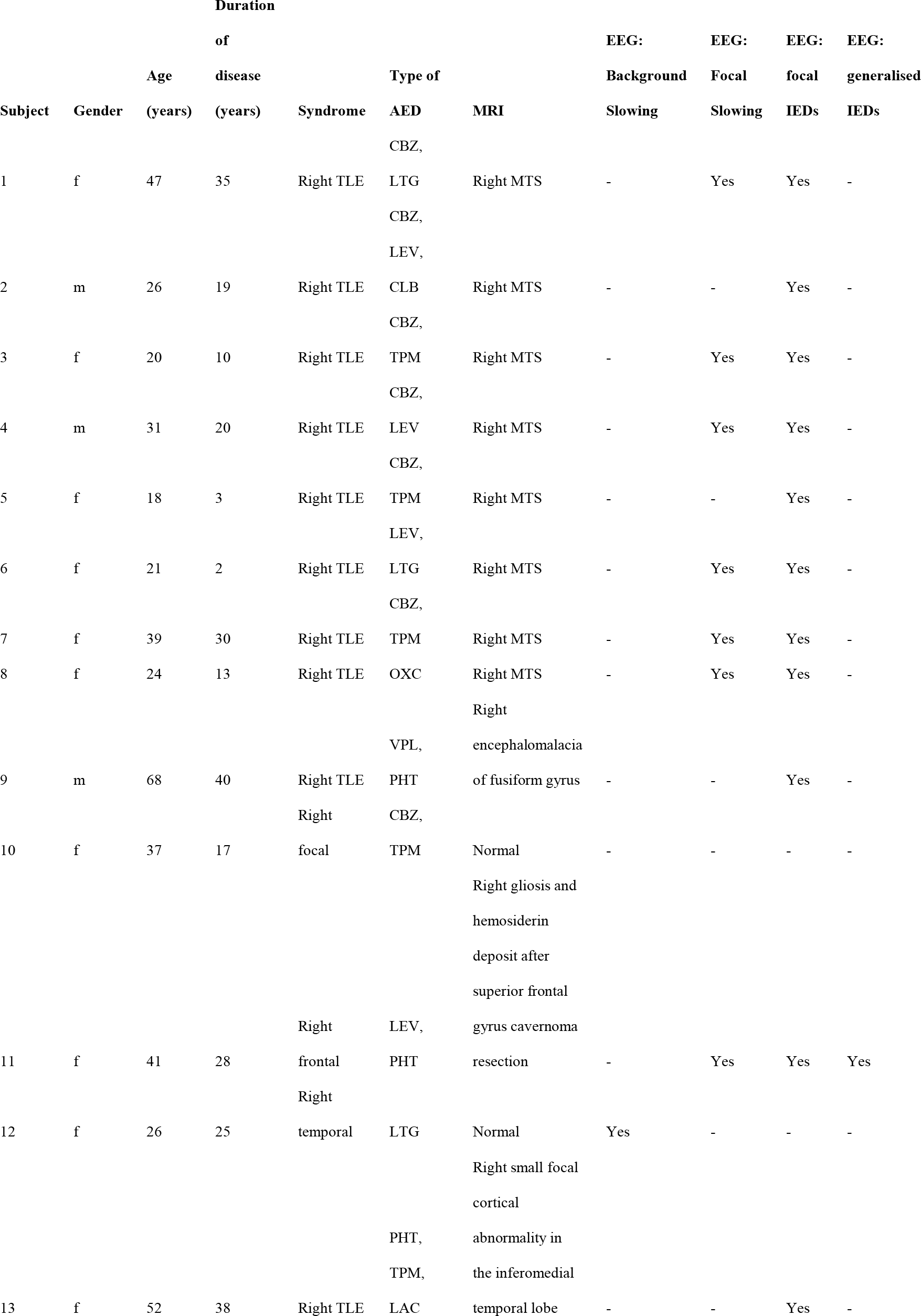
Right Focal Epilepsy Cohort.

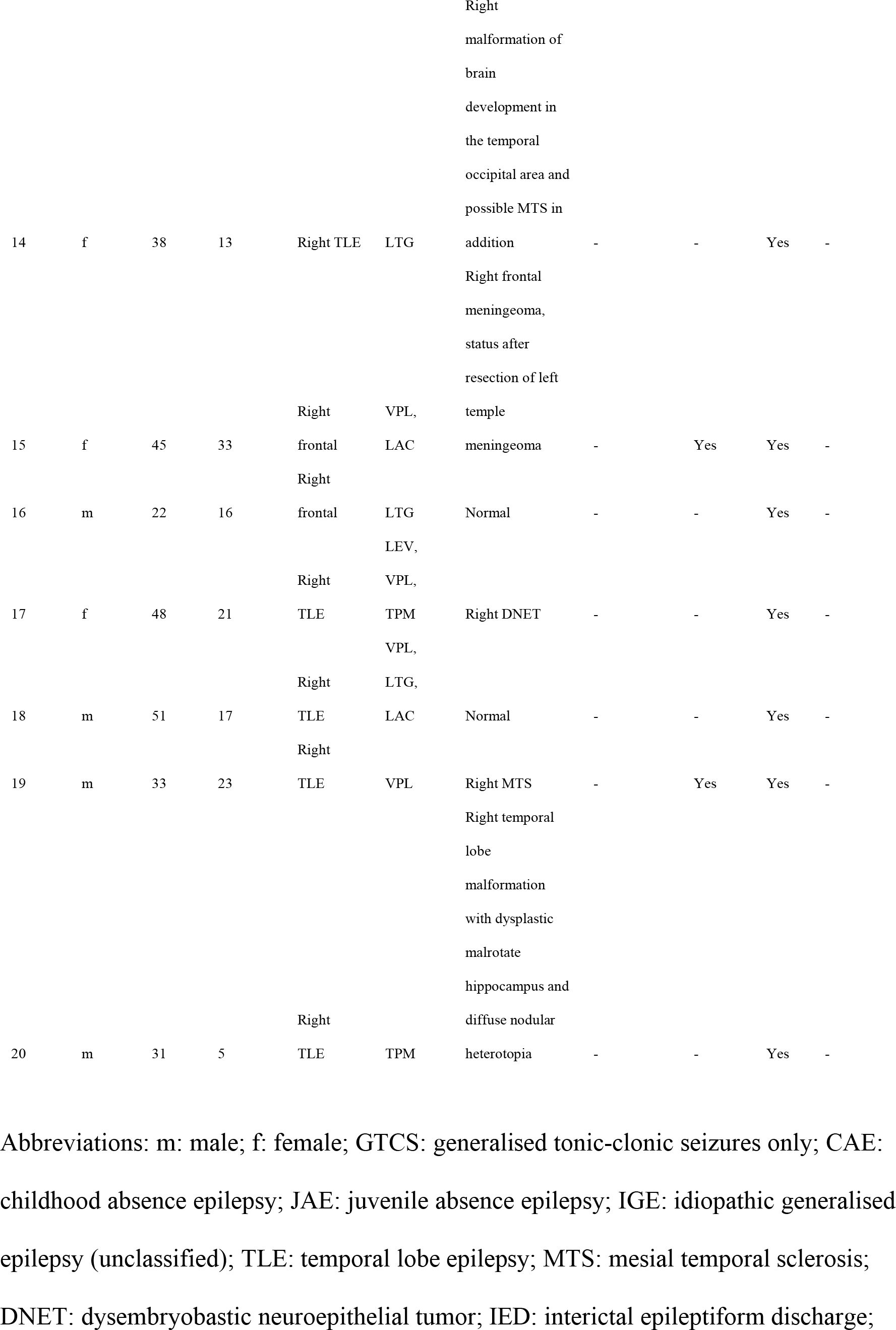

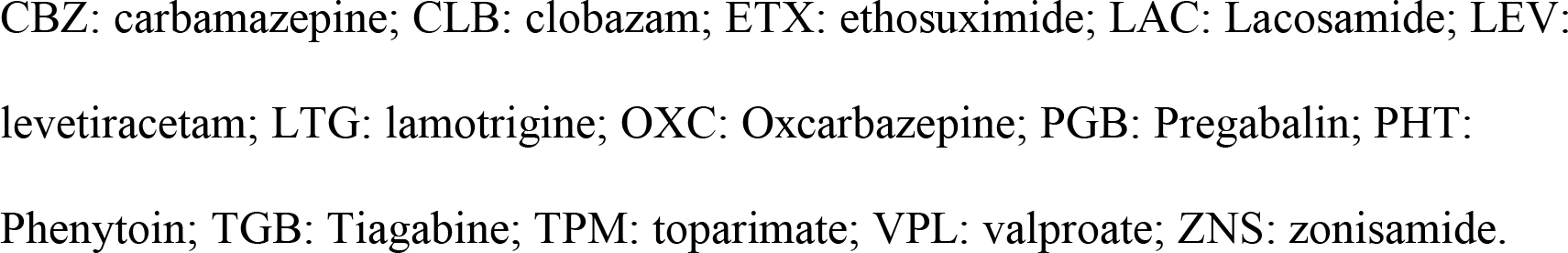

Table: Group Comparisons.

**Table: Group Comparisons**
|  | Sex | Age | Duration | Number of AEDs |
| --- | --- | --- | --- | --- |
| **Controls – IGE** | 0.4341 | 0.4991 | - | - |
| **Controls – Left Focal** | 0.4053 | 0.1065 | - | - |
| **Controls – Right Focal** | 0.4398 | 0.1043 | - | - |
| **IGE – Left Focal** | 0.1482 | 0.4436 | 0.2075 | 0.4479 |
| **IGE – Right Focal** | 0.9999 | 0.3686 | 0.4786 | 0.1533 |
| **Left Focal – Right Focal** | 0.1292 | 0.7607 | 0.0902 | 0.4643 |

*Caption: The Mann-Whitney U test (two-tailed, $p < 0.05$ ) was used to compare age, duration of epilepsy and drug load between the groups. Fisher's exact test ( $p < 0.05$ ) was used for comparing sex. All tests failed to reject the null-hypothesis of equal medians of the groups.*

## References

1. Fornito A, Zalesky A, Breakspear M. The connectomics of brain disorders. Nat Rev Neurosci. 2015;16:159–172.

2. de Tisi J, Bell GS, Peacock JL, et al. The long-term outcome of adult epilepsy surgery, patterns of seizure remission, and relapse: a cohort study. The Lancet 2011;378:1388–1395.

3. Keller SS, Richardson MP, Schoene-Bake JC, et al. Thalamotemporal Alteration and Postoperative Seizures in Temporal Lobe Epilepsy. Ann Neurol. 2015;77:760–774.

4. Goodfellow M, Rummel C, Abela E, Richardson MP, Schindler KA, Terry JR. Estimation of brain network ictogenicity predicts outcome from epilepsy surgery. Nat Sci Reports. 2016;6:29215.

5. Goodfellow M, Rummel C, Abela E, Richardson MP, Schindler K, Terry JR. Computer models to inform epilepsy surgery strategies: prediction of postoperative outcome. Brain 2017;140:e30.

6. Proix T, Bartolomei F, Guye M, Jirsa VK. Individual brain structure and modelling predict seizure propagation. Brain 2017;140:641–654.

7. Sinha N, Dauwels J, Kaiser M, et al. Predicting neurosurgical outcomes in focal epilepsy patients using computational modelling. Brain 2016;140:319–332.

8. Meeren HKM, Pijn JPM, van Luijtelaar ELJM, Coenen AML, Lopes Da Silva FH. Cortical Focus Drives Widespread Corticothalamic Networks during Spontaneous Absence Seizures in Rats. J Neurosci. 2002;22:1480–1495.

9. Terry JR, Benjamin O, Richardson MP. Seizure generation: The role of nodes and networks. Epilepsia 2012;53:e166–169.

10. Fisher RS, Cross JH, French JA, et al. Operational classification of seizure types by the International League Against Epilepsy: Position Paper of the ILAE Commission for Classification and Terminology. Epilepsia 2017;58:522–530.

11. Scheffer IE, Berkovic S, Capovilla G, et al. ILAE classification of the epilepsies: position paper of the ILAE Commission for Classification and Terminology. Epilepsia 2017;58:512–521.

12. Schmidt H, Petkov G, Richardson MP, Terry JR. Dynamics on Networks: The Role of Local Dynamics and Global Networks on the Emergence of Hypersynchronous Neural Activity. PLoS Comput Biol. 2014;10:e1003947.

13. Schmidt H, Woldman W, Goodfellow M, et al. A computational biomarker of idiopathic generalized epilepsy from resting state EEG. Epilepsia 2016;57:e200–204.

14. Chowdhury FA, Woldman W, FitzGerald TH, et al. Revealing a brain network endophenotype in families with idiopathic generalised epilepsy. PloS one 2014;9:e110136.

15. Pampiglione G, Kerridge J. E.E.G. abnormalities from the temporal lobe studied with sphenoidal electrodes. J Neurol Neurosurg Psychiatry 1956;19:117–129.

16. Gudmundsson S, Runarsson TP, Sigurdsson S, Eiriksdottir G, Johnsen K. Reliability of quantitative EEG features. Clinical Neurophysiology 2007;118(10):2162–2171.

17. Fraschini M, Demuru M, Crobe A, Marrosu F, Stam CJ, Hillebrand A. The effect of epoch length on estimated EEG functional connectivity and brain network organisation. Journal of Neural Engineering 2016;13(3),036015.

18. Shackman AJ, McMenamin BW, Maxwell JS, Greischar LL, Davidson RJ. Identifying robust and sensitive frequency bands for interrogating neural oscillations. Neuroimage 2010;51:1319–1333.

19. Larsson PG, Kostov, H. Lower frequency variability in the alpha activity in EEG among patients with epilepsy. Clinical Neurophysiology 2005;116:2701–2706.

20. Lachaux J, Rodriguez E, Martinerie J, Varela FJ. Measuring Phase Synchrony in Brain Signals. Hum Brain Mapp. 1999;8:194–208.

21. Stam CJ, Nolte G, Daffertshofer A. Phase lag index: Assessment of Functional Connectivity From Multi Channel EEG and MEG With Diminished Bias from Common Sources. Hum Brain Mapp. 2007;28:1178–1193.

22. Schreiber T, Schmitz A. Improved Surrogate Data for Nonlinearity Tests. Phys Rev Lett. 1996;77:635–638.

23. Kuramoto Y. Cooperative Dynamics of Oscillator Community. Prog Theor Phys Suppl. 1984;79:223–240.

24. Schuster HG, Wanger P. A model for neuronal oscillators in the visual cortex. Biol Cybern. 1990;64:77–82.

25. Breakspear M, Heitmann S, Daffertshofer A. Generative models of cortical oscillations: neurobiological implications of the Kuramoto model. Front Hum Neurosci. 2010;4:1–14.

26. Daffertshofer A, van Wijk BCM. On the influence of amplitude on the connectivity between phases. Front Neuroinform. 2011;5:1–12.

27. Hamandi K, Salek-Haddadi A, Laufs H, et al. EEG-fMRI of idiopathic and secondarily generalized epilepsies. Neuroimage 2006;31:1700–1710.

28. Berg AT, Berkovic SF, Brodie MJ, et al. Revised terminology and concepts for organization of seizures and epilepsies: report of the ILAE Commission on Classification and Terminology, 2005-2009. Epilepsia 2010;51:676–685.

29. Smith SJM. EEG in the diagnosis, classification, and management of patients with epilepsy. J Neurol Neurosurg Psychiatry 2005;76(Suppl II):ii2–ii7.

30. Benbadis SR, Lin K. Errors in EEG Interpretation and Misdiagnosis of Epilepsy: Which EEG Patterns Are Overread? Eur Neurol. 2008;59:267–271.

31. Smith D, Defalla BA, Chadwick DW. The misdiagnosis of epilepsy and the management of refractory epilepsy in a specialist clinic. Q J Med. 1999;92:15–23.

32. Chowdhury FA, Nashef L, Elwes RDC. Misdiagnosis in epilepsy: a review and recognition of diagnostic uncertainty. Eur J Neurol. 2008;15:1034–1042.

33. van Diessen E, Otte WM, Stam CJ, Braun KPJ, Jansen FE. Electroencephalography based functional networks in newly diagnosed childhood epilepsies. Clin Neurophysiol. 2016;127:2325–2332.

34. Englot DJ, Hinkley LB, Kort NS, et al. Global and regional functional connectivity maps of neural oscillations in focal epilepsy. Brain 2015;138:2249–2262.

35. Pedersen M, Omidvarnia AH, Walz JM, Jackson GD. Increased segregation of brain networks in focal epilepsy: An fMRI graph theory finding. NeuroImage Clin. 2015;8:536–542.

36. Verhoeven T, Coito A, Plomp G, et al. Automated diagnosis of temporal lobe epilepsy in the absence of interictal spikes. NeuroImage: Clinical 2018;17:10–15.

37. Badawy RAB, Freestone DR, Lai A, Cook MJ. Epilepsy: Ever-changing states of cortical excitability. Neuroscience 2012;222:89–99.

